# Analysis of potential mechanisms of non-carbapenemase mediated carbapenem resistance in *Acinetobacter baumannii*

**DOI:** 10.1101/2025.01.13.632675

**Authors:** Lisa-Marie Höfken, Sören G. Gatermann, Niels Pfennigwerth

## Abstract

**Objectives:** *Acinetobacter baumannii* is a Gram-negative nosocomial pathogen that plays an important role in the context of bacterial multidrug-resistance. Increasing resistance to carbapenems in particular is of high therapeutic relevance, as in this case only a few antibiotics remain for treatment. The mechanisms of carbapenem resistance in *A. baumannii* are mainly based upon carbapenemases and the mechanisms such as porin loss, efflux pumps and altered PBPs have been poorly studied to date.

**Methods:** The *A. baumannii* reference strain ATCC 17978 was artificially mutated by selection pressure with increasing meropenem concentrations until carbapenem resistance was achieved. Growth analyses were carried out with the mutants and MICs for relevant antibiotics were determined. In addition, the mutants were whole genome sequenced, and the sequences were compared with the wild type. As various mutagenesis attempts for targeted construction of these respective mutants were unfortunately not successful, the strain collection of the NRC was screened for isolates that showed carbapenem resistance without a detectable carbapenemase. These isolates were sequenced and analysed for abnormalities in PBPs and porins in comparison to the sequence of the reference strain ATCC 17978 and were compared to other strains that possessed a carbapenemase.

**Results:** In three of the resulting ATCC 17978 mutants, a mutation of PBP2 was observed (W366L). Theses mutants were carbapenem resistant and were not affected by avibactam in contrast to the wild type ATCC 17978. Growth experiments indicated a fitness loss compared to the wild type.

As W366L was not found in the clinical isolates, we looked for other abnormalities in various genes associated with carbapenem resistance. Mutations were primarily found in the PBPs, with a mutation in PBP3 (A515V) occurring particularly often.

**Conclusion:** The results of this work support the prevailing thesis that PBP mutations in *A. baumannii* can lead to carbapenem resistance. Since there are hardly any studies on this hypothesis and for the most part only using outdated methods, these results are of particular relevance and further studies on this topic are recommended.

## Introduction

*Acinetobacter baumannii* is a nonfermentative, Gram-negative, nonmotile and oxidase-negative bacillus. It can be found commonly in soil and water. Because of its environmental resistance, particularly to dryness, it has also been able to spread easily in the environment of hospitals and healthcare facilities. It is a very effective human colonizer in the hospital. It has been shown that care workers hands are frequently colonized with *Acinetobacter* species, resulting in further spread and nosocomial infections.^1, 2^ *Acinetobacter baumannii* was susceptible to most antibiotics until the 1970s, but since then it rapidly acquired many resistance determinants to a wide range of antibacterial agents.^2^ This is a serious and ongoing problem in the treatment of infections. However, attempts to limit the spread of rapidly evolving antibiotic-resistant pathogens are best achieved by a detailed understanding of the causes that drive these resistance patterns.

### Resistance mechanisms

For *Acinetobacter* there are already many resistance mechanisms known. Resistance to carbapenems, which are used as last resort antibiotics against *Acinetobacter* infections is of clear therapeutic relevance, as it leaves only polymyxins and tigecycline available for treatment in many cases. The most common reason for carbapenem resistance in *Acinetobacter baumannii* is the presence of a carbapenemase. In Germany, the most frequently found are OXA-23, OXA-72 and NDM-1.^3^ But there are also other mechanisms, that confer resistance to carbapenems. CarO, a 29-kDa outer membrane channel protein which confers resistance to both imipenem and meropenem in *Acinetobacter* has been well characterised.^4–6^ OprD is a well-studied porin frequently associated with imipenem resistance in *Pseudomonas aeruginosa* and a homologue of OprD was found in *Acinetobacter*, suggesting the same resistance mechanism.^7^ Furthermore, reduced production of 37-, 44-, and 47 kDa OMPs in carbapenem-resistant isolates together with increased production of class C cephalosporinase has been shown to lead to carbapenem resistance.^8^ Overexpression of AdeABC efflux pump may also confer high-level resistance to carbapenems in conjunction with carbapenem-hydrolysing oxacillinases.^9^ However, little is known about the role of PBPs in carbapenem resistance in *Acinetobacter* to date.^10^

### Aim of this work

The German National Reference Centre for multidrug-resistant Gram-negative bacteria (NRC) has a huge collection of over 80,000 resistant pathogens. In this collection, there are several carbapenem-resistant *Acinetobacter baumannii* isolates, most of them possessing a carbapenemase gene. However, a small portion of isolates are carbapenem-resistant without producing a carbapenemase and without any other known reason for carbapenem resistance. The aim of this work was to investigate possible reasons for non-carbapenemase-mediated carbapenem resistance in *A. baumannii*.

## Material/Methods

### Bacterial strains

All isolates analysed in this study were clinical isolates from patients hospitalised in various German hospitals that were sent to the NRC for routine surveillance purposes. 75 % of the isolates originated from infections processes. Species identification was performed by using MALDI-TOF-MS (Bruker Daltonics, Billerica, MA, USA).

All tested strains and information on origin, sequence type and β-lactamases are listed in Table 2.

### Carbapenemase detection

For phenotypic characterisation and for the detection or exclusion of carbapenemases, the *Acinetobacter baumannii* isolates were examined using a modified hodge test and a synergy test with EDTA, combined with PCRs on the carbapenemase enconding genes *bla*_OXA-23_, *bla*_OXA-24/40_, *bla*_OXA-58_, *bla*_NDM_, followed by Sanger sequencing of PCR products.^11–13^

### MICs

MICs were determined in triplicate using broth microdilution (BMD) with pre-configured microtiter plates (Bruker, Bremen, Germany) with CAMHB and evaluated according to EUCAST breakpoints.^14^

### Selection of spontaneous mutants of *A. baumannii* ATCC 17978

*A. baumannii* ATCC 17978 was plated on blood agar plates and incubated overnight at 37 °C. Next step was a macrodilution with meropenem with concentrations of up to 128 mg/L in 4 ml LB-media. Briefly, ATCC 17978 was dispensed in 0.1 % NaCl to a McFarland of 0.5 and each vial of the macrodilution was inoculated with 50 µl of the suspension. After incubation at 37 °C overnight at 200 rpm, cells of the vial with the highest concentration of meropenem where growth was visible were taken to inoculate the next macrodilution as described before. In some cases, these cells took two days to grow in the presence of meropenem. When the cells did not reach a higher level of meropenem after 3-5 cycles of incubation, a cryo-culture of this cells was prepared.

### Growth experiments

Determination of relative cell fitness was done via monoculture growth curve experiments. Biomass monitoring was performed in triplicate using a Cell Growth Quantifier (CGQ; aquila biolabs GmbH, Baesweiler, Germany). Overnight cultures were inoculated in 50 mL of LB-media to an OD_600 nm_ of 0.05 ± 0.005. Shaking flasks were put on an optical sensor array and incubated for 24 h at 150 rpm and 37 °C. Data were evaluated using the CGQuant software (aquila biolabs GmbH, Baesweiler, Germany).

### Sequencing

The molecular detection of carbapenemase genes was carried out by PCRs, as described previously (Primers used are shown in Table S1).^15^ PCR products were purified using the NucleoSpin® Gel and PCR Clean-up kit (Macherey-Nagel, Düren, Germany) and then Sanger-sequenced. If the PCR did not yield any results, the isolates were subjected to Whole-genome sequencing. Genomic Tips Kit (Qiagen, Hilden, Germany) was used to purify high molecular weight DNA. Libraries were created with Nextera® XT Library Prep Kit (Illumina, Eindhoven, The Netherlands). The sequencing was performed *in-house* on a MiSeq2 platform with 2 × 300 bp paired-end reads (Illumina, Eindhoven, The Netherlands). The resulting fastq-files were assembled using Spades version 3.13.1. An *in-house* gene search tool based on a database consisting of numerous genes associated with β-lactam resistance in *A. baumannii* received from NCBI was used for screening. Present ORFs were checked for correctness using BLAST (https://blast.ncbi.nlm.nih.gov/) and were annotated using RAST (Rapid Annotations using Subsystems Technology).^16–18^

### Genetic characterisation of spontaneous mutants

The WGS sequencing data of the *A. baumannii* ATCC 17978 and its spontaneous mutants were checked for SNPs via snippy on the Galaxy server (Galaxy Version 4.6.0+galaxy0).^19^

### Mutagenesis studies

Several mutagenesis techniques were tried to create PBP mutations in ATCC 17978 chromosomally. Among others, the CRSIPR-Cas-9 system of Wang et al,^20^ homologous recombination mediated by the RecAB recombination system, deletion of a PBP via resistance cassettes and suicide vectors were used.^21^

## Results

Three of the subpopulations of ATCC 17978 survived up to a meropenem concentration of 32 mg/L. These three putative mutants of ATCC 17978 were subjected to whole-genome-sequencing together with the original ATCC 17978 to compare the sequences and to screen for putative mutations that increased meropenem resistance.

Surprisingly, all three mutants showed only one single SNP compared to the origin ATCC 17978, which was located in the *mrd*A gene, resulting in the amino acid substitution W366L in PBP2.

MICs of the *A. baumannii* ATCC 17978 PBP2 mutants were determined, and the results showed an increase in MICs of imipenem from 0.125 mg/L for the wild type and 2 to 4 mg/L for the mutants (Table 2). A large increase in MICs of meropenem and ertapenem could also be seen (from 0.25 to 32 mg/L and 1 to 64 mg/L for meropenem and ertapenem, respectively). MICs for aztreonam increased by only one dilution step higher from 32 to 64 mg/L, but for the wildtype the combination with avibactam lowered the MIC to 8 mg/L, whereas avibactam had no effect on the mutants. MICs for ceftozolan-avibactam were higher in the mutants than in the wildtype and increased ≤ 0.25 to 8 mg/L. MICs for ceftazidim-avibactam, colistin and tigecyclin were not affected by the W366L mutation of PBP2.

Growth experiments with the wild type *A. baumannii* ATCC 17978 and the selected PBP2 mutants showed that ATCC 17978 grew faster and achieved an overall higher cell density (OD_600_ of 3.7) than the PBP2 mutants (OD_600_ of 2.8) (figure 1).

It was planned to verify the effect of the PBP2 mutations by mutagenesis studies in the sensitive reference strain, but unfortunately approaches with various mutagenesis techniques were not successful due to failing homologous recombination in ATCC 17978.

Instead, all *Acinetobacter baumannii* isolates that were received by the NRC since 2009 that were tested meropenem-resistant but did not harbour a carbapenemase (n = 18) were tested again with phenotypical tests and PCRs to exclude any potentially overseen carbapenemases and subjected to whole genome sequencing to screen for the mutations observed in the selection experiments.

A carbapenemase could not be detected in any of the following isolates: NRZ-18984, NRZ-29926, NRZ-30058, NRZ-41034, NRZ-44141, NRZ-52866, NRZ-53861, NRZ-56398, NRZ-69687 and NRZ-86385 (further information on origin and sequence type in table 1). The MICs of these isolates reached 0,25 to 2 mg/L for imipenem, 2 to 4 mg/L for meropenem and 8 to 16 mg/L for ertapenem (Table 3).

**Table 1.** MICs of *A. baumannii* ATCC 17978 and its mutants in mg/L.

| Antibiotic | ATCC 17978 WT | ATCC 17978 mutant 1 | ATCC 17978 mutant 2 | ATCC 17978 mutant 3 |
| --- | --- | --- | --- | --- |
| Imipenem | 0.125 | 4 | 4 | 2 |
| Imipenem-Relebactam<sup>c</sup> | ≤ 0.25 | 4 | 4 | 2 |
| Meropenem | 0.25 | 32 | 32 | 16 |
| Meropenem-Vaborbactam<sup>d</sup> | ≤ 1 | 32 | 32 | 16 |
| Ertapenem | 1 | 64 | 64 | 64 |
| Aztreonam | 32 | 64 | 64 | 64 |
| Aztreonam-Avibactam<sup>a</sup> | 8 | 64 | 64 | 64 |
| Ceftazidim-Avibactam<sup>a</sup> | 16 | 16 | 16 | 16 |
| Ceftozolan-Tazobactam<sup>b</sup> | ≤ 0.25 | 8 | 8 | 8 |
| Colistin | 2 | 2 | 2 | 2 |
| Tigecyclin | ≤ 0.125 | ≤ 0.125 | ≤ 0.125 | ≤ 0.125 |

<sup>a</sup> Fixed concentration of Avibactam of 4 mg/L
<sup>b</sup> Fixed concentration of Tazobactam of 4 mg/L
<sup>c</sup> Fixed concentration of Relebactam of 4 mg/L
<sup>d</sup> Fixed concentration of Vaborbactam of 8 mg/L

**Table 2.** Background information about all tested isolates in this study, including origin, sequence type and β-lactamases.

| Strains without carbapenemase | Type of isolate | location | Year of isolation | Sequence type (if available) | | | β-lactamases |
| --- | --- | --- | --- | --- | --- | --- | --- |
|  |  |  |  | MLST Oxford | Pasteur | cgMLSTv1 |  |
| NRZ-18984 | clinical isolate | rectal swab (stool) | 2015 | 440 | 2 | cgST-1663 | OXA-64*, GES-12, ADC-25 |
| NRZ-29926 | clinical isolate | oral pharyngeal swab | 2016 | 1806, 208 | 2 | cgST-2249 | OXA-66*, TEM-1, ADC-25 |
| NRZ-30058 | clinical isolate | bronchial lavage | 2016 | 1806, 208 | 2 | cgST-4409 | OXA-66*, ADC-25 |
| NRZ-41034 | clinical isolate | rectal swab (stool) | 2018 |  |  | cgST-1662 | OXA-66*, ADC-25 |
| NRZ-44141 | clinical isolate | tracheal secretion | 2018 | 1806, 208 | 2 | cgST-4200 | OXA-66*, TEM-1, ADC-25 |
| NRZ-52866 | clinical isolate | unknown | 2019 | 2614 | 1724 |  | OXA-203*, ADC-25 |
| NRZ-56961 | clinical isolate | rectal swab (stool) | 2019 | 1806,208 | 2 | cgST-5840 | OXA-66*, TEM-1, ADC-25 |
| NRZ-69687 | clinical isolate | tracheal cannula swab | 2021 | 1806,208 | 2 | cgST-5854 | OXA-66*, TEM-1, ADC-25 |
| NRZ-86385 | clinical isolate | tracheal secretion | 2023 | 1806, 208 | 2 | cgST-6734/-7186 | OXA-66*, TEM-1, ADC-25 |
| Strains with carbapenemase |  |  |  |  |  |  |  |
| NRZ-41108 | clinical isolate | bronchial lavage | 2018 | 1324 | 374 |  | OXA-825, OXA-259*, ADC-25 |
| NRZ-74350 | clinical isolate | oral pharyngeal swab | 2022 | 1806, 208 | 2 | cgST-4144 | OXA-72, OXA-66*, ADC-25 |
| NRZ-75439 | clinical isolate | wound swab | 2022 | 1806, 208 | 2 | cgST-742 | OXA-23, OXA-66*, ADC-25 |
| NRZ-76416 | clinical isolate | rectal swab (stool) | 2022 | 1100 | 400 |  | GES-11, OXA-100*, ADC-25 |
| NRZ-79748 | clinical isolate | rectal swab (stool) | 2022 | 218 | 570 |  | NDM-1, OXA-23, OXA-66*, TEM-1, ADC-25 |
| NRZ-80289 | clinical isolate | rectal swab (stool) | 2022 | 2839 | 78 |  | OXA-72, OXA-90*, CARB-14, CTX-M-124, ADC-25 |
| NRZ-80805 | clinical isolate | punctate | 2022 | 1816, 195 | 2 | cgST-3690 | OXA-23, OXA-66*, ADC-25 |
| NRZ-80930 | clinical isolate | blood culture | 2022 |  |  | cgST-1744/-1747/-48 | OXA-72, OXA-90*, CARB-14, ADC-25 |
| NRZ-83921 | clinical isolate | wound swab | 2023 | 942 | 267 |  | KPC-23, OXA-180*, TEM-1, ADC-25 |
| NRZ-86003 | clinical isolate | wound swab | 2023 | 1816,195 | 2 | cgST-1298 | OXA-23, OXA-66*, ADC-25 |
| NRZ-87256 | clinical isolate | oral pharyngeal swab | 2023 | 450 | 2 |  | PER-7, OXA-23, OXA-66*, ADC-25 |
| NRZ-88408 | clinical isolate | swab groin | 2023 | 2062,2063 | 2 | cgST-7247 | PER-7, OXA-23, OXA-66*, ADC-25 |

\* all betalactamases that are marked with an \* belong to the OXA-51-like family

**Table 3.** MICs of *Acinetobacter baumannii* isolates in mg/L.

| Antibiotic | NRZ-18984 | NRZ-29926 | NRZ-30058 | NRZ-41034 | NRZ-44141 | NRZ-52866 | NRZ-53861 | NRZ-56398 | NRZ-69687 | NRZ-86385 |
| --- | --- | --- | --- | --- | --- | --- | --- | --- | --- | --- |
| Imipenem | 4 | 2 | 2 | 2 | 0.25 | 2 | 2 | 2 | 2 | 0.5 |
| Imipenem-Relebactam<sup>c</sup> | 4 | 2 | 1 | 1 | ≤ 0.25 | 1 | 1 | 1 | 1 | 0.5 |
| Meropenem | 8 | 4 | 4 | 4 | 4 | 2 | 2 | 4 | 2 | 8 |
| Meropenem-Vaborbactam<sup>d</sup> | 8 | 4 | 4 | 4 | 4 | 2 | 2 | 2 | 2 | 8 |
| Ertapenem | 16 | 16 | 16 | 16 | 8 | 16 | 8 | 16 | 8 | 16 |
| Aztreonam | > 64 | 64 | 64 | > 64 | > 64 | 64 | > 64 | > 64 | > 64 | > 64 |
| Aztreonam-Avibactam<sup>a</sup> | > 64 | 32 | 64 | > 64 | > 64 | 64 | 64 | > 64 | 64 | > 64 |
| Ceftazidim-Avibactam<sup>a</sup> | 32 | > 32 | 32 | > 32 | > 32 | > 32 | 32 | > 32 | 32 | > 32 |
| Ceftozolan-Tazobactam<sup>b</sup> | > 8 | > 8 | > 8 | > 8 | > 8 | > 8 | > 8 | > 8 | > 8 | > 8 |
| Colistin | 2 | 2 | 2 | 2 | 2 | 2 | 2 | 1 | 2 | 1 |
| Tigecyclin | 1 | 0.5 | 2 | 1 | 0.5 | 1 | 1 | 1 | 1 | 0.5 |

<sup>a</sup> Fixed concentration of Avibactam of 4 mg/L
<sup>b</sup> Fixed concentration of Tazobactam of 4 mg/L
<sup>c</sup> Fixed concentration of Relebactam of 4 mg/L
<sup>d</sup> Fixed concentration of Vaborbactam of 8 mg/L

**Table 4.**
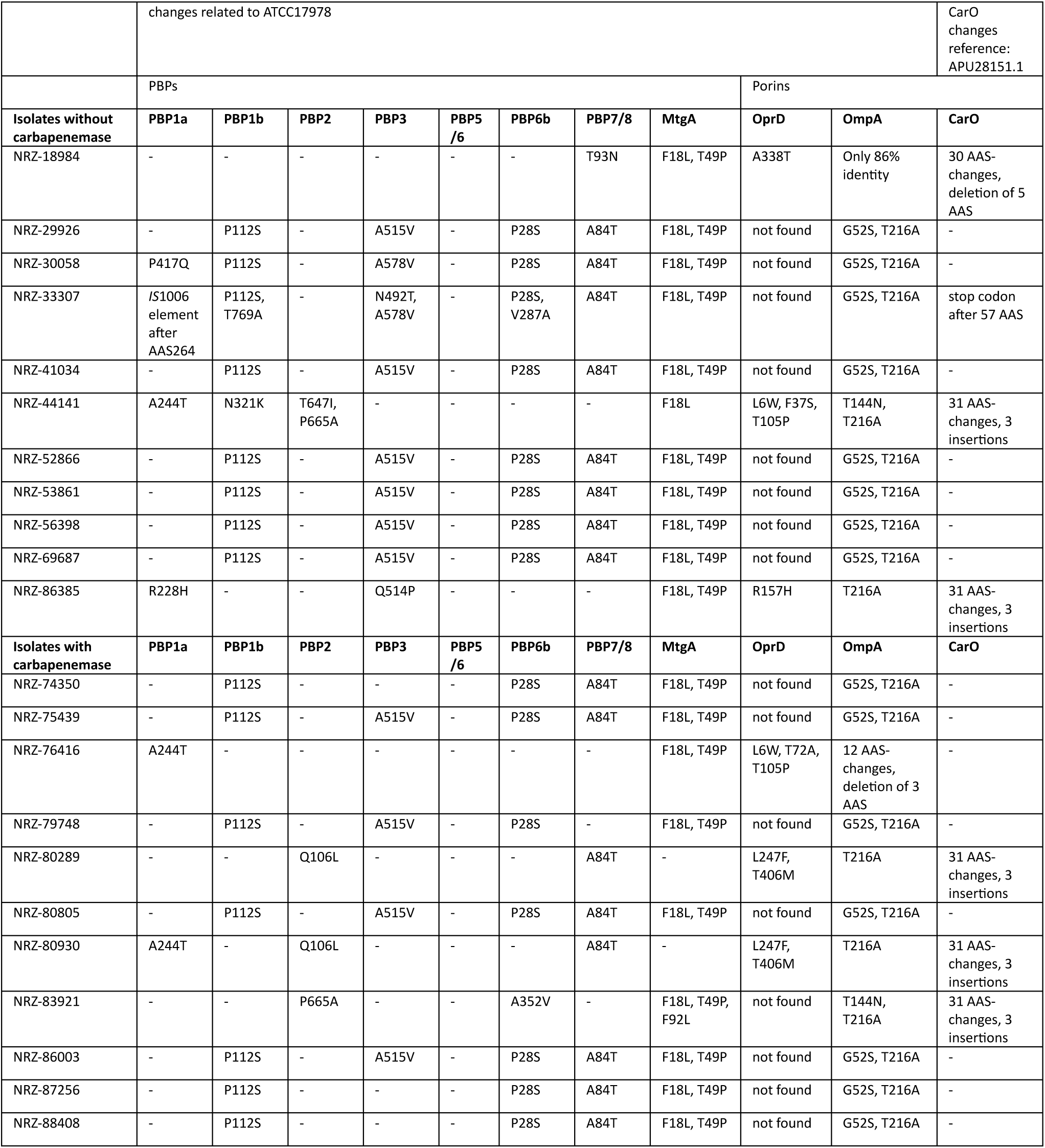
Results of the sequence comparison of selected proteins of the *A. baumannii* strains with and without carbapenemase in comparison to the reference strain ATCC 17978, or in the case of CarO to APU28151.1, as the reference strain has no intact CarO. (AAS=amino acids)

These 11 isolates were subjected to whole genome sequencing via Illumina to check if there are also PBP mutations that were found in the *in vitro* experiment with ATCC 17978. 11 other *Acinetobacter baumannii* isolates that were carbapenem-resistant and carried a carbapenemase (Table 1) were also sequenced for comparison.

Isolates without a carbapenemase had the following mutations: A244T in PBP1a was found in two strains, P417Q and R228H in PBP1a in one isolate each and a disruption of *mrc*A after 264 amino acids due to an *IS*1006 element was found in one isolate. The other 7 isolates had an identical PBP1a to ATCC 17978. Seven isolates showed an amino acid change in PBP1b, P112S, and one isolates with this amino acid change and additional T769A. One isolate had an amino acid change at another position in the protein, N321K. For PBP2 of the isolates without carbapenemase two amino acid changes occurred in only one isolate; T647L and P665A. PBP3 showed more often amino acid changes, there could be found seven times the amino acid change A515V, one-time N492T together with A578V and one-time Q514P. For PBP5/6 all isolates showed the same amino acid sequence as ATCC 17978. Eight isolates showed the PBP6b amino acid change P28S compared to ATCC 17978 and in one isolate occurred V287A. The protein MtgA showed ten times the amino acid change F18L and T49P and one-time F18L alone.

The isolates with a carbapenemase had the following amino acid changes compared to ATCC 17978. Only two times a mutated PBP1a was found with the amino acid change A244T. Seven isolates had also the amino acid changes P112S in PBP1b. Two strains showed the mutations Q106L and P665A in PBP2. None of these strains had the W366L mutation. A515V in PBP3 was also found four times. For PBP5/6 all isolates showed the same amino acid sequence as ATCC 17978. For PBP6b the amino acid change P28S occurred in seven isolates and in one isolate there was the amino acid change A352V. For MtgA the isolates showed eight times F18L together with T49P and one-time additional F92L.

The porins OprD, CarO and OmpA, that are typically relevant for carbapenem resistance in *Acinetobacter baumannii*, were also checked for these isolates. CarO was not found in ATCC 17978, so another reference sequence was used (APU28151.1).

Sequence comparison of this porins with the isolates without carbapenemase resulted for OprD in eight times in no results, in one isolate there was the amino acid change A388T, in another L6W, F37S and T105P and in the last one R175H. CarO showed in two isolates 31 amino acid changes together with three insertions and one time 30 amino acid changes with five deleted amino acids. In one isolate there could only be found a protein with 86% identity to OmpA, eight isolates showed G52S and T216A in OmpA, one isolate T144N with T216A and one isolate only T216A.

Results for the isolates with a carbapenemase were similar, in eight isolates it was not possible to find OprD, in two isolates there were OprD proteins with L247F and T406M and in one isolate there were three amino acid changes: L6W, T72A and T105P. CarO showed in three isolates 31 amino acid changes together with three insertions. OmpA of these isolates showed seven times G52S with T216A, two times only T216A alone, one time T144n with T216A and for one isolate 12 amino acid changes together with a deletion of three amino acids were found for OmpA.

## Discussion

Although it has already been described in detail for many bacterial species that altered PBPs can lead to carbapenem resistance,^22^ there are very few studies on this for *Acinetobacter baumannii*. Some of them are over 20 years old and the methodology was limited at the time.

In 1991, Gehrlein and colleagues analysed the *A. baumannii* strain 4852/88 and some imipenem-resistant clones of the same strain. They saw an alteration in the PBPs in an SDS-PAGE, but it was not possible to say which PBPs were affected in detail.^23^

12 years later, in 2003, a study was published by Fernandez-Cuenca *et al*., in which various *A. baumannii* isolates with different resistance phenotypes were investigated using iodine-125 conjugate-labelled PBP binding assays.^24, 25^ The results were visualised by autoradiography in an SDS-PAGE and showed different patterns of PBPs. They concluded that the absence of a band which they assumed to be to PBP2, leads to carbapenem resistance. However, this conclusion is problematic as they only named the PBPs according to size, i.e. the largest PBP was PBP1 and so on, which is contradictory to PBP nomenclature in other species.

In 2011, a paper was published that for the first time provided an overview of the PBPs in *A. baumannii* in relation to the genetic background.^26^ It was reported that when compared to the PBPs of *E. coli*, *A. baumannii* has four high molecular weight (PBP1a, PBP1b, PBP2 and PBP3) and 3 low molecular weight PBPs (PBP5/6, PBP6b and PBP7/8) as well as the monofunctional MtgA (figure 2).

However, the estimated sizes of the PBPs of *A. baumannii* ATCC 17978 do not correlate with the numbering that is used in *E. coli*, which significantly complicates literature reviews. PBP1a is not the largest PBP with 55.1 kDa, PBP1b is bigger with 88.2 kDa. Therefore, if in older publications PBPs were separated by SDS gels according to their size and then the PBPs were simply numbered from the largest to the smallest, this is incorrect with regard to the correlation of naming in *E. coli*. This must be considered when quoting specific statements on PBPs from the corresponding publications. Statements that altered PBP profiles could have an effect on carbapenem resistance are of course still valid.

But it has to be considered that many carbapenemases were not yet known at that time and therefore may simply not have been found in some strains, as there was no whole genome sequencing performed to check this more precisely.

In the abovementioned study, Cayo *et al*. also analysed several *A. baumannii* isolates from hospitals, with some of them being imipenem-sensitive and others resistant (in relation to the ECOFF). The PBP-encoding genes were sequenced and several mutations found, but they were present in both the sensitive and the resistant strains. However, all resistant isolates carried an OXA-24 carbapenemase or had an insertion of IS*Aba1* upstream of *bla*_OXA-51-like_ and/or the AmpC-encoding gene, which is known to result in carbapenem resistance. Furthermore, the altered PBPs may only lead to increased antibiotic tolerance, which does not necessarily exceed the ECOFF, so that isolates with those PBP mutations may still have been categorised as susceptible.

In 2011, Vashist *et al*. published a study in which the PBPs of 20 ß-lactam-resistant strains were compared with the PBPs of ATCC 17978.^27^ This was done by tagging the PBPs with Bocillin FL and subsequent size separation in an SDS-PAGE. Different band patterns and thus different compositions of PBPs were found, but again the previously mentioned problem of correctly labelling the PBPs based on size alone was encountered. Moreover, smaller point mutations or similar small changes cannot be recognised in an SDS gel.

PBP2 from *A. baumannii* found to have a zinc binding site in the transpeptidase domain.^28^ Zinc appears to be critical for the stability of PBP2 and to maintain a normal cell wall shape. Mutations that interfere with the zinc binding site have led to a loss of function of PBP2 and increased ß-lactam resistance. The amino acids D350, D365, H371 and C384 are probably responsible for the binding of zinc. The mutation W366L in PBP2 in our spontaneous mutants may also have these effects especially because the base exchange is directly one amino acid next to the potential zinc-binding amino acid D365.

It was furthermore shown that mutations in AdeB (F136L and G280S) and PBP3, including A515V, lead to meropenem MICs of 32 mg/L and that A515V is very close to the binding site for meropenem and therefore probably influences binding. ^29^ In this study, the A515V mutation in PBP3 was also found frequently in *A. baumannii* isolates, that were carbapenem resistant without harbouring a carbapenemase. These results support the hypothesis of Hawkey *et al*. that this particular mutation could lead to meropenem resistance in *A. baumannii*.

A mutagenesis study on the D,D-transpeptidases of *A. baumannii* and discovered that the PBP3 of *A. baumannii* could not be deleted and is therefore essential for the growth and secondly, mutants lacking the three PBPs 1a, 1b and 2 were up to 8 times more sensitive to ß-lactams.^30^

Not much has been published on the involvement of PBPs in carbapenem resistance in *Acinetobacter* spp., but since PBPs do play a role in other species, it is reasonable to assume that this is also the case here. The results of this study show the potential involvement of an *in vitro* induced PBP2 mutation in carbapenem resistance. In some carbapenem resistant isolates from our strain collection, in which no carbapenemase could be detected, some PBP mutations were also found, mainly A515V in PBP3. As this mutation was also found in other strains with carbapenemase, further studies would be interesting to see whether carbapenem resistance is also prevalent without a carbapenemase present.

Mutagenesis studies were attempted in this study, but were unfortunately unsuccessful, as *A. baumannii* ATCC 17978 did not allow homologous recombination, regardless of the mutagenesis technique (CRISPR-Cas9, assisted homologous recombination, suicide vectors or deletion cassettes). Sometimes initial screening gave the impression of successful cloning, but after sequencing it turned out that the relevant resistance cassettes had only been incorporated into transposons or IS elements and that no homologous recombination had taken place at the desired site. For this reason, various mutagenesis techniques were tried and varied, but all failed due to the missing homologous recombination.

A slightly reduced growth of the mutants compared to the wild type strain possible indicates a loss of fitness due to the PBP2 mutation. This could explain why such mutations are rarely found in wild *Acinetobacter baumannii* isolates. However, there is still a risk that such a mutation could occur during carbapenem therapy and lead to treatment failure without horizontal gene transfer and the involvement of carbapenemases. In combination with some β-lactamases or porin losses or efflux pumps, PBP mutations can have some important effects on the resistance profile.

The role of Omps in antibiotic resistance is very speculative because most *Acinetobacter* Omps haven’t even been annotated or studied yet. However, there is some research on OmpA, OprD and CarO and their role in β-lactam resistance.

OmpA is the most prevalent porin in *A. baumannii* and disruption mutants were found to have increased susceptibility to aztreonam, chloramphenicol, imipenem, meropenem and nalidixic acid. ^31^ Studies also suggest that OmpA may be coupled to efflux pumps that force antimicrobial compounds out of the periplasmic space. ^32^

Most of the *A. baumannii* isolates examined in this study had only two amino acid differences compared to ATCC 17978 in OmpA (G52S, T216A). Consequently, this could simply be an allelic variation. Mutagenesis and expression studies would be required to further investigate the influence on β-lactam resistance. However, there is one strain that does not appear to have intact OmpA (NRZ-18984), which could potentially lower its resistance level against carbapenems compared to the other isolates.

CarO is another *A. baumannii* porin associated with carbapenem susceptibility. Some studies observed a selective imipenem influx in *A. baumannii*, ^5^ while another study created a liposome model system embedded with CarO that denied the ability to transport carbapenems through CarO. ^33^ Despite this solitary observation, there is a wealth of evidence from different research groups pointing to the role of CarO in antibiotic resistance. ^34^

In this study there were several isolates with a wild-type CarO, but also some isolates with an altered form with 30 or more amino acid substitutions. This may also contribute to carbapenem resistance, independent of the presence of a carbapenemase.

This study also found that the sensitive reference strain ATCC 17978 does not possess CarO, which, according to the current state of research, could lead to imipenem resistance. However, this strain is absolutely sensitive to imipenem. The loss of CarO alone does not appear to be sufficient to confer relevant resistance to imipenem. However, in combination with other resistance determinants, it may have an additive effect.

OprD was first identified in studies of the outer membrane of carbapenem-resistant *A. baumannii* isolates. ^7^ It is an orthologue of a porin involved in the transport of basic amino acids and imipenem in *P. aeruginosa.* ^35^ While an isogenic deletion mutant of *A. baumannii* OprD did not affect the MICs of β-lactams, ^32^ a significant reduction in the MIC of imipenem, ertapenem and meropenem was observed in another *Acinetobacter* species, *A. baylyi* spp.. ^36^ Despite these contradictory reports of OprD and antibiotic resistance in *A. baumannii*, SNPs and insertion elements in OprD were frequently identified in MDR A. baumannii, which may indicate a role for OprD in resistance. In this study about three quarters of isolates did not pocess the oprD porin and all other isolates showed single amino acid changes in this porin.

No correlation could be drawn between abnormalities in porins and the presence of a carbapenemase in the isolates examined here. Further studies are needed to break down the complex influence of porins on the resistance of *Acinetobacter baumannii*.

## Conclusion

Many reviews on *Acinetobacter baumannii* state that carbapenem resistance is caused by PBPs, among other mechanisms. However, a closer look at the studies available to date reveals that the data on this is rather sparse and should be viewed with caution. In this study, it was found that resistance to meropenem could be induced *in vitro* with a sensitive reference strain, resulting from a single amino acid exchange in PBP2. In addition, further PBP mutations were found in other clinical isolates that were carbapenem-resistant but do not possess a carbapenemase, further underlining the potential importance of PBP mutations for carbapenem resistance. Consequently, further studies on this topic are essential.

Furthermore, several mutations in the porins OprD, CarO or OmpA were found in the strains analysed in this study. More studies on the influence of porins in *A. baumannii* on the resistance to antimicrobial compounds are also urgently needed, especially since many porins have not yet been described in this species.

**Figure 1.**
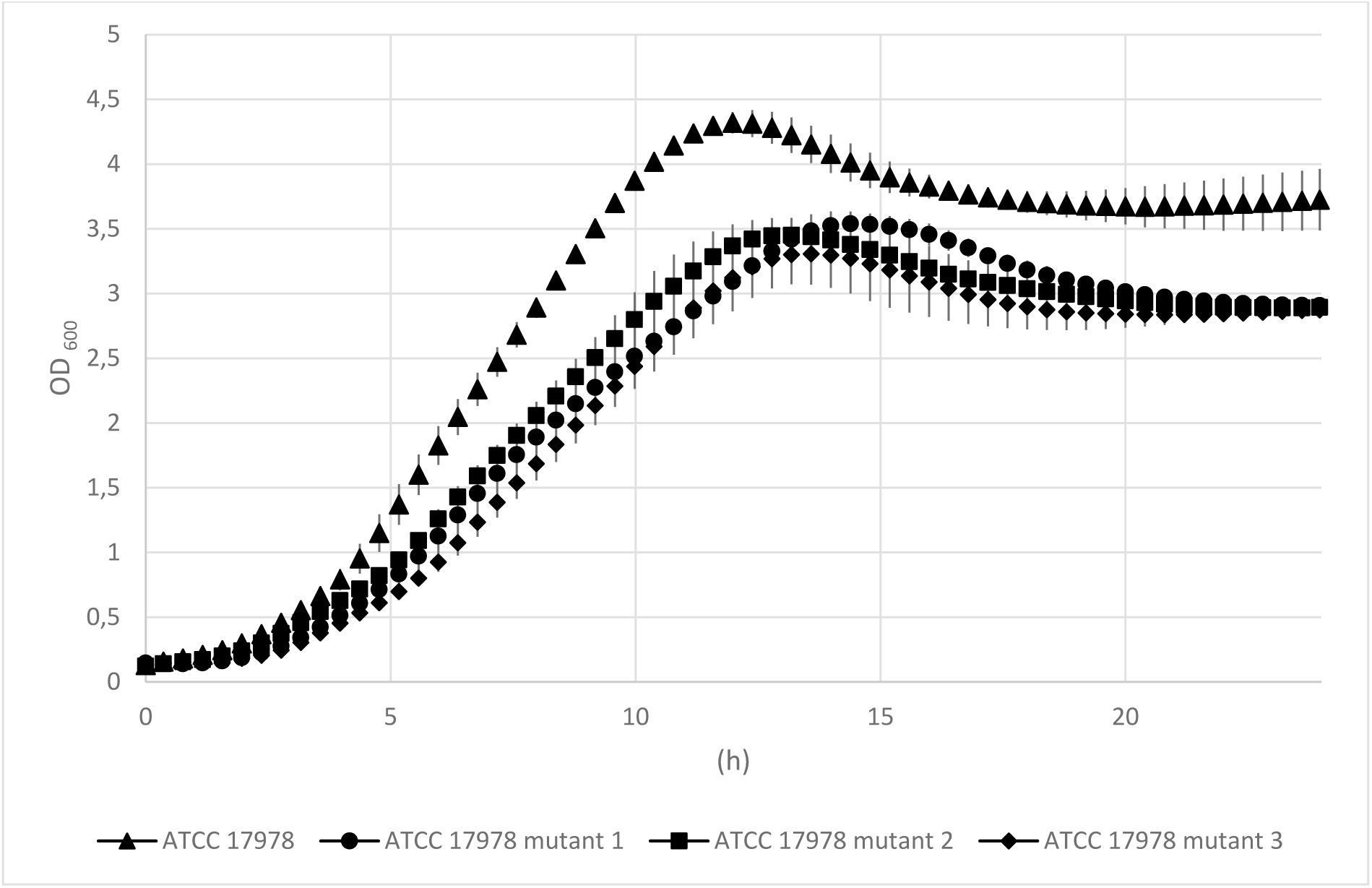
Growth experiment of *A. baumannii* ATCC 17978 and its mutants with PBP2 mutation. Measurements were performed in 50 ml LB medium at 37°C for 24 h, all cultures were inoculated to an OD_600_ of 0.05. The mean value and the standard deviation from three growth experiments are shown. Measurements were made using CGQ.

**Figure 2.**
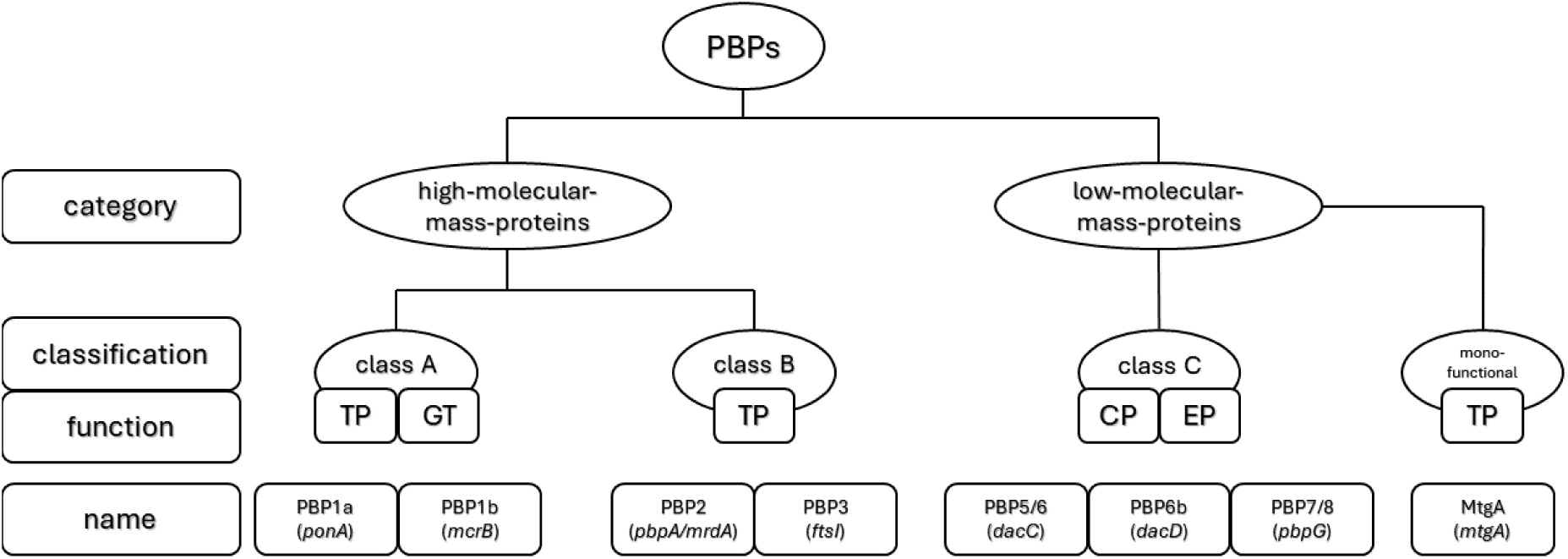
Overview on PBPs of *Acinetobacter baumannii* Figure adapted from Lange *et al*., 2019 (TP – transpeptidase, GT – glycosylase, CP – carboxypeptidase, EP – endopeptidase

## Acknowledgements

The authors thank Anke Albrecht, Nadine Frey, Susanne Friedrich, Brigitte Hemmerle, Svenja Hirle, Anja Kaminski, Kirsten Krengel, Ulrike Maduch, Lea Rogge, Marion Schmidt, Laura Suppa, and Joanna Waniczek for excellent technical assistance.

## Transparency declarations

S.G.G. has received speaker fees from Beckman Coulter and bioMérieux.

N.P. has received speaker or consultancy fees from bioMérieux, Pfizer and Shionogi. All other authors: none to declare.

## References

1. Almasaudi SB. *Acinetobacter spp*. as nosocomial pathogens: Epidemiology and resistance features. Saudi journal of biological sciences 2018; 25: 586–96.

2. Lee K, Yong D, Jeong SH, et al. Multidrug-Resistant *Acinetobacter spp*.: Increasingly Problematic Nosocomial Pathogens. Yonsei medical journal 2011; 52: 879.

3. Pfennigwerth N. Tätigkeitsbericht des Nationalen Referenzzentrums für gramnegative Krankenhauserreger für den Zeitraum 01. Januar 2023 bis 31. Dezember 2023.

4. Limansky AS, Mussi MA, Viale AM. Loss of a 29-kilodalton outer membrane protein in *Acinetobacter baumannii* is associated with imipenem resistance. Journal of clinical microbiology 2002; 40: 4776–8.

5. Mussi MA, Limansky AS, Viale AM. Acquisition of resistance to carbapenems in multidrug-resistant clinical strains of *Acinetobacter baumannii*: natural insertional inactivation of a gene encoding a member of a novel family of beta-barrel outer membrane proteins. Antimicrobial agents and chemotherapy 2005; 49: 1432–40.

6. Siroy A, Molle V, Lemaître-Guillier C et al. Channel formation by CarO, the carbapenem resistance-associated outer membrane protein of *Acinetobacter baumannii*. Antimicrobial agents and chemotherapy 2005; 49: 4876–83.

7. Dupont M, Pagès J-M, Lafitte D, et al. Identification of an OprD homologue in *Acinetobacter baumannii*. Journal of proteome research 2005; 4: 2386–90.

8. Quale J, Bratu S, Landman D et al. Molecular epidemiology and mechanisms of carbapenem resistance in *Acinetobacter baumannii* endemic in New York City. Clinical Infectious Diseases: An Official Publication of the Infectious Diseases Society of America 2003; 37: 214–20.

9. Marqué S, Poirel L, Héritier C et al. Regional occurrence of plasmid-mediated carbapenem-hydrolyzing oxacillinase OXA-58 in *Acinetobacter spp*. in Europe. Journal of clinical microbiology 2005; 43: 4885–8.

10. Roca I, Espinal P, Vila-Farrés X et al. The *Acinetobacter baumannii* Oxymoron: Commensal Hospital Dweller Turned Pan-Drug-Resistant Menace. Frontiers in microbiology 2012; 3: 148.

11. Nordmann P, Poirel L, Carrër A et al. How To Detect NDM-1 Producers. Journal of Clinical Microbiology 2020; 49: 718–21.

12. Woodford N, Ellington Mj Fau - Coelho JM, Coelho Jm Fau - Turton JF et al. Multiplex PCR for genes encoding prevalent OXA carbapenemases in *Acinetobacter spp*.. International Journal of Antimicrobial Agents 2006; 27: 10–1016

13. Seyedi M, Yousefi F, Naeimi B, et al. Phenotypic and genotypic investigation of metallo-β-lactamases in *Pseudomonas aeruginosa* clinical isolates in Bushehr, Iran. . Iran J Basic Med Sci. 2022; 25: 1196–1200

14. Testing TECoAS. Breakpoint tables for interpretation of MICs and zone diameters, version 14.0. http://www.eucast.org/clinical_breakpoints/.

15. Pfennigwerth N, Gatermann SG, Körber-Irrgang B et al. Phenotypic Detection and Differentiation of Carbapenemase Classes Including OXA-48-Like Enzymes in Enterobacterales and *Pseudomonas aeruginosa* by a Highly Specialized Micronaut-S Microdilution Assay. Journal of clinical microbiology 2020; 58.

16. Altschul SF, Gish W, Miller W et al. Basic local alignment search tool. Journal of molecular biology 1990; 215: 403–10.

17. Camacho C, Coulouris G, Avagyan V et al. BLAST+: architecture and applications. BMC bioinformatics 2009; 10: 421.

18. Aziz RK, Bartels D, Best AA et al. The RAST Server: Rapid Annotations using Subsystems Technology. BMC Genomics 2008; 9: 75.

19. The Galaxy C. The Galaxy platform for accessible, reproducible, and collaborative data analyses: 2024 update. Nucleic Acids Research 2024; 52: W83–W94.

20. Wang Y, Wang Z, Ji Q. CRISPR-Cas9-Based Genome Editing and Cytidine Base Editing in *Acinetobacter baumannii*. STAR Protocols 2020; 1: 100025.

21. Schäfer A, Tauch A, Jäger W et al. Small mobilizable multi-purpose cloning vectors derived from the *Escherichia coli* plasmids pK18 and pK19: selection of defined deletions in the chromosome of *Corynebacterium glutamicum*. Gene 1994; 145: 69–73.

22. Sauvage E, Kerff F, Terrak M, et al. The penicillin-binding proteins: structure and role in peptidoglycan biosynthesis. FEMS microbiology reviews 2008; 32: 234–58.

23. Gehrlein M, Leying H, Cullmann W et al. Imipenem resistance in *Acinetobacter baumanii* is due to altered penicillin-binding proteins. Chemotherapy 1991; 37: 405–12.

24. Cuenca FF, Pascual A, Martínez Marínez L et al. Evaluation of SDS-polyacrylamide gel systems for the study of outer membrane protein profiles of clinical strains of *Acinetobacter baumannii*. Journal of basic microbiology 2003; 43: 194–201.

25. Fernández-Cuenca F, Martínez-Martínez L, Conejo MC et al. Relationship between beta-lactamase production, outer membrane protein and penicillin-binding protein profiles on the activity of carbapenems against clinical isolates of *Acinetobacter baumannii*. The Journal of antimicrobial chemotherapy 2003; 51: 565–74.

26. Cayô R, Rodríguez M-C, Espinal P et al. Analysis of genes encoding penicillin-binding proteins in clinical isolates of *Acinetobacter baumannii*. Antimicrobial agents and chemotherapy 2011; 55: 5907–13.

27. Vashist J, Tiwari V, Das R et al. Analysis of penicillin-binding proteins (PBPs) in carbapenem resistant *Acinetobacter baumannii*. The Indian Journal of Medical Research 2011; 133: 332–8.

28. Micelli C, Dai Y, Raustad N et al. A conserved zinc-binding site in *Acinetobacter baumannii* PBP2 required for elongasome-directed bacterial cell shape. Proceedings of the National Academy of Sciences of the United States of America 2023; 120: e2215237120.

29. Hawkey J, Ascher DB, Judd LM et al. Evolution of carbapenem resistance in *Acinetobacter baumannii* during a prolonged infection. Microbial genomics 2018; 4.

30. Toth M, Lee M, Stewart NK et al. Effects of Inactivation of d,d-Transpeptidases of *Acinetobacter baumannii* on Bacterial Growth and Susceptibility to β-Lactam Antibiotics. Antimicrobial agents and chemotherapy 2022; 66: e0172921.

31. Kwon HI, Kim S, Oh MH et al. Outer membrane protein A contributes to antimicrobial resistance of *Acinetobacter baumannii* through the OmpA-like domain. Journal of Antimicrobial Chemotherapy 2017; 72: 3012–5.

32. Smani Y, Fàbrega A, Roca I et al. Role of OmpA in the multidrug resistance phenotype of *Acinetobacter baumannii*. Antimicrob Agents Chemother 2014; 58: 1806–8.

33. Zahn M, D’Agostino T, Eren E et al. Small-Molecule Transport by CarO, an Abundant Eight-Stranded β-Barrel Outer Membrane Protein from *Acinetobacter baumannii*. Journal of molecular biology 2015; 427: 2329–39.

34. Uppalapati SR, Sett A, Pathania R. The Outer Membrane Proteins OmpA, CarO, and OprD of *Acinetobacter baumannii* Confer a Two-Pronged Defense in Facilitating Its Success as a Potent Human Pathogen. Frontiers in Microbiology 2020; 11.

35. Hancock RE, Brinkman FS. Function of pseudomonas porins in uptake and efflux. Annu Rev Microbiol 2002; 56: 17–38.

36. Morán-Barrio J, Cameranesi MM, Relling V et al. The Acinetobacter Outer Membrane Contains Multiple Specific Channels for Carbapenem β-Lactams as Revealed by Kinetic Characterization Analyses of Imipenem Permeation into *Acinetobacter baylyi* Cells. Antimicrob Agents Chemother 2017; 61.

