## Supplementary Table 1 for "Analysis of potential mechanisms of non-carbapenemase mediated carbapenem resistance in *Acinetobacter baumannii*"

**Table S1** Primers used in this study to screen for *bla* genes.

| Gene | Sequence (5'-≥3') | Annealing Temperature(°C) | Amplicon size (bp) | Reference |
| --- | --- | --- | --- | --- |
| *bla*_KPC_ | TGTCACTGTATCGCCGTC | 58 | 1009 | ^37^ |
|  | CTCAGTGCTCTACAGAAAACC |  |  |  |
| *bla*_VIM_ | GTTTGGTCGCATATCGCAAC | 54 | 382 | ^38^ |
|  | AATGCGCAGCACCAGGATAG |  |  |  |
| *bla*_IMP_ | GAAGGYGTTTATGTTCATAC | 54 | 587 | ^38^ |
|  | GTAMGTTTCAAGAGTGATGC |  |  |  |
| *bla*_OXA-48_ | TTGGTGGCATCGATTATCGG | 52 | 744 | ^39^ |
|  | GAGCACTTCTTTTGTGATGGC |  |  |  |
| *bla*_GIM_ | AGTTGATTCGAATGGGTTGGT | 55 | 748 | NRC^a^ |
|  | TCGCCTACGTAACCTAAGCC |  |  |  |
| *bla*_SPM_ | CCTACAATCTAACGGCGACC | 55 | 629 | ^40^ |
|  | TCGCCGTGTCCAGGTATAAC |  |  |  |
| *bla*_DIM_ | GTACCTGAGCTAAGAATCGAG | 50 | 668 | NRC^a^ |
|  | CGGCTGGATTGATTTGTTAGAG |  |  |  |
| *bla*_NDM_ | TGCCCAATATTATGCACCCG | 55 | 780 | NRC^a^ |
|  | CAGCGCAGCTTGTCGG |  |  |  |
| *bla*_AIM_ | GAAACGTCGCTTCACCCTG | 50 | 482 | NRC^a^ |
|  | ACCAGGATGTCGCAGTCGAG |  |  |  |
| *bla*_KHM_ | GCTCTTGTTATATCGTTTGGTC | 55 | 626 | NRC^a^ |
|  | CATTGTTGCATTGCTATAACGG |  |  |  |
| *bla*_SMB_ | CAGCAGCCATTCACCATCTA | 50-55 | 491 | ^41^ |
|  | GAAGACCACGTCCTTGCACT |  |  |  |
| *bla*_TMB_ | GGATTGGAAGTTGAGGAAATTGAC | 50 | 476 | NRC^a^ |
|  | GCACCACAATCTTAGCTTCAG |  |  |  |
| *bla*_FIM_ | GAAGCACATGGAAAACTGGG | 50-55 | 431 | ^42^ |
|  | GATGGGCGAATGAGACAGC |  |  |  |
| *bla*_HMB_ | CATTGGGTCTTTTGCTGCTG | 50-55 | 687 | NRC^a^ |
|  | CCGCCTCAAGCGCAC |  |  |  |
| *bla*_CAM_ | CGTTTTCGGGCAAACGGGC | 55 | 633 | NRC^a^ |
|  | GATTTCGTGCTTGCCCATCCGTC |  |  |  |

^a^in-house primers designed for the routine diagnostics of the NRC.

**1**	Yigit H, Queenan AM, Anderson GJ *et al.* Novel carbapenem-hydrolyzing b-lactamase, KPC-1, from a carbapenem-resistant strain of *Klebsiella pneumoniae*. *Antimicrob Agents Chemother* 2001; **45**: 1151-61.

**2**	Pitout JD, Gregson DB, Poirel L *et al.* Detection of *Pseudomonas aeruginosa* producing metallo-b-lactamases in a large centralized laboratory. *J Clin Microbiol* 2005; **43**: 3129-35.

**3**	Poirel L, Héritier C, Tolün V *et al.* Emergence of oxacillinase-mediated resistance to imipenem in *Klebsiella pneumoniae*. *Antimicrob Agents Chemother* 2004; **48**: 15-22.

**4**	Castanheira M, Toleman MA, Jones RN *et al.* Molecular characterization of a b-lactamase gene, *bla*_GIM-1_, encoding a new subclass of metallo-b-lactamase. *Antimicrob Agents Chemother* 2004; **48**: 4654-61.

**5**	Wachino J, Yoshida H, Yamane K *et al.* SMB-1, a novel subclass B3 metallo-b-lactamase, associated with I*SCR1* and a class 1 integron, from a carbapenem-resistant *Serratia marcescens* clinical isolate. *Antimicrob Agents Chemother* 2011; **55**: 5143-9.

**6**	Pollini S, Maradei S, Pecile P *et al.* FIM-1, a new acquired metallo-β-lactamase from a *Pseudomonas aeruginosa* clinical isolate from Italy. *Antimicrob Agents Chemother* 2013; **57**: 410-6.
